# Integrative Genome-wide Analysis of the Determinants of RNA Splicing in Kidney Renal Clear Cell Carcinoma

**DOI:** 10.1101/010256

**Authors:** Kjong-Van Lehmann, André Kahles, Cyriac Kandoth, William Lee, Nikolaus Schultz, Oliver Stegle, Gunnar Rätsch

**Author notes:** Both authors contributed equally (Kjong-Van Lehmann, André Kahles).

## Abstract

We present a genome-wide analysis of splicing patterns of 282 kidney renal clear cell carcinoma patients in which we integrate data from whole-exome sequencing of tumor and normal samples, RNA-seq and copy number variation. We proposed a scoring mechanism to compare splicing patterns in tumor samples to normal samples in order to rank and detect tumor-specific isoforms that have a potential for new biomarkers. We identified a subset of genes that show introns only observable in tumor but not in normal samples, ENCODE and GEUVADIS samples. In order to improve our understanding of the underlying genetic mechanisms of splicing variation we performed a large-scale association analysis to find links between somatic or germline variants with alternative splicing events. We identified 915 *cis*- and *trans*-splicing quantitative trait loci (sQTL) associated with changes in splicing patterns. Some of these sQTL have previously been associated with being susceptibility loci for cancer and other diseases. Our analysis also allowed us to identify the function of several COSMIC variants showing significant association with changes in alternative splicing. This demonstrates the potential significance of variants affecting alternative splicing events and yields insights into the mechanisms related to an array of disease phenotypes.

## 1. Introduction

The analysis of gene expression and the identification of expression quantitative trait loci (eQTLs) has become a standard part of the analyses performed in many population genetics studies. However, the variability in expression levels is only one of the factors shaping the complexity of the transcriptome. RNA-modifying processes, especially the process of alternative splicing, enable the formation of several RNA isoforms from a single gene locus and drastically increase transcriptome complexity. During splicing, specific parts are excised from the pre-mRNA (introns) and the remaining parts (exons) are re-connected. Through combinatorial choice of introns, different mRNAs can be generated. This tightly regulated process is also termed alternative splicing.^1,2^ The role of alternative splicing in cancer is being actively investigated,^3,4^ however it is often difficult to separate tissue specific effects from tumor specific changes. Defects in the splicing machinery or dysregulation of the process can lead to disease or play an active role in cancer progression.^5–7^ Interestingly, several strategies involving natural compounds or antisense oligonucleotides have been suggested to target aberrant splicing,^8,9^ making the detection of alternative splicing events as drug targets desirable. Yet, splicing efficiency has only recently been considered as a quantitative trait in genetic analysis. First studies describing systematically alternative splicing in the context for genomic variation have been conducted by Battle *et al.*^10^ and in context of the GEUVADIS project.^11^ Also studies with a specific focus on *single* alterations that affect splicing have been published recently, for instance, the identification of somatic mutations in U2AF1 causative for altered splicing in acute myeloid leukemias^12^ as well as identification of a somatic variant affecting SF3B1 function.^13^

In this work, we present the genome-wide analysis of alternative splicing events in 282 Kidney Renal Clear Cell Carcinoma (KIRC) samples generated in context of The Cancer Genome Atlas project (TCGA).^14^ We perform an integrative analysis of RNA-seq, whole-exome and copy number variation, in order to identify determinants of splicing variation caused by germline and somatic genetic variation. We first built a comprehensive inventory of alternative splicing events occurring in KIRC tumors and characterized tumor-specific splicing controlling for tissue specific effects. Encouraged by the presence of cancer-specific introns, we used a mixed model approach to systematically associate splicing alterations with germline and somatic genetic variants with the aim to identify splicing quantitative trait loci (sQTLs). This analysis enables us to shed some light on genetic mechanisms underlying alternative splicing patterns in cancer and normal cells.

## 2. Methods

We here provide an outline of the methodology taken. Please note that a detailed description of our methods can be found in the supplemental material.

### 2.1 Data Processing

Matching 282 whole exome and transcriptome samples have been downloaded from cgHub and were realigned using STAR. For comparison purposes we have also downloaded and realigned 140 RNA-Seq samples from the GEUVADIS project as well as 460 RNA-Seq samples from the ENCODE project. Expression counts have been generated based on the GENCODE annotation and splicing phenotypes have been generated using SplAdder.^15^

Germline variants have been called using the HaplotypeCaller in GATK and somatic variants have been identified using MuTect.

### 2.2 Tumor specific splicing analysis

Tumor-specific splicing has been identified by ranking all expressed genes by the ratio of the average number of samples that expressed a certain intron in the KIRC tumor samples over the average number of samples expressing the intron in KIRC normals, GEUVADIS and ENCODE combined. Functional enrichment analysis has been undertaken by making use of the GOrilla webserver.^16^

### 2.3 Quantitative Trait Analysis

Quantification of splicing measured in PSI has been performed using an inverse normal transform resolving ties randomly and variants have been encoded numerically under an additive genetic model (see Supplemental methods for details). A linear mixed model analysis has been used to find associations between germline mutations and splicing changes. We accounted for population structure as well as possible hidden confounders using PANAMA and known confounders as in gene expression and copy number variation from Ciriello *et al.*^17^. Associations have been computed using LIMIX^18^ and Benjamini-Hochberg step-up procedure has been used for FDR estimation to correct for multiple testing.

## 3. Results

### 3.1 Identification of Tumor-specific Splicing

Based on the splicing graph constructed with SplAdder, we extracted 184, 941 introns located in 15, 387 genes that were part of alternative splicing events. Of these introns, 160, 208 were confirmed by at least 10 spliced alignments in at least one of the samples from the KIRC, GEUVADIS or ENCODE sets. Interestingly, when ranked by exclusive occurrence in tumor samples (see methods), especially transmembrane proteins of the solute carrier family (SLC) comprising a family of roughly 450 genes, were significantly enriched amongst the top ranks showing a 12 fold enrichment (p-value 3.6 *·* 10^−5^, hypergeom. test; compare Fig. 1, Panel A). Although single members of this family have been related to cancer biology, e.g., SLC28A1,^19^ in general not much is known about their function in context of cancer. Other top-ranked membrane-associated proteins show stronger known links to cancer, such as the transmembrane collagen COL23A1^20^ or the transmembrane protein 176A.^21^ Encouragingly, the latter as well as its heterologous protein TMEM176B also appeared on top of the list of genes that show exclusive intron expression and harbor significant sQTL (Fig. 1, Panel B). It is notable that the accumulation of TMEM176A/B has been linked to several other cancer types^21^ previously. Moreover, we also found non-membrane associated genes with exclusive intron expression that are known to have links to cancer, such as secretagogin (SCGN)^22^ involved in cell proliferation, the cytochrome P450 epoxygenase CYP2J2^23^ or the hypoxia-inducible factor EGLN3.^24^ We observed that most exclusive introns were indeed result of splicing and not an artifact of lacking gene expression, although several genes show considerably less expression in normal samples. (see Supplemental Figure 1).

**Fig. 1.**
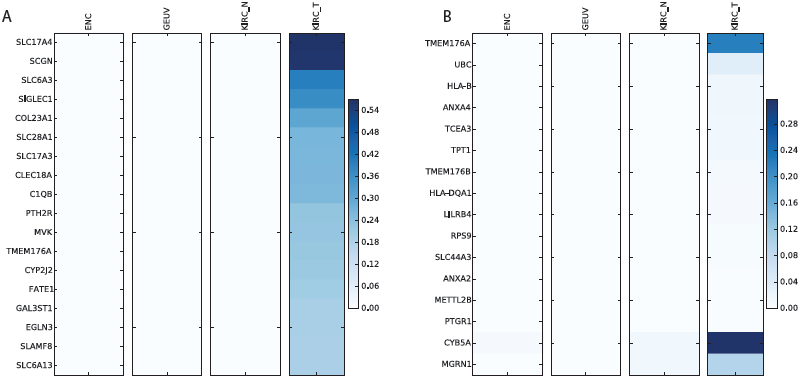
Enrichment of introns exclusively present in tumor samples. *A:* List of genes that contain the top 20 most exclusive introns. Color represents fraction of samples that have confirmed this intron with 10 alignments. In all cases we observe no or little evidence of these introns in the control samples. *B:* List of genes with an sQTL that contain exclusively expressed introns. Color represents fraction of samples having confirmed the intron with *≥* 10 alignments.

In agreement to these findings, a functional enrichment analysis on gene ontology (GO) categories showed significant enrichment of membrane transport processes but also in extracellular matrix organization and amino-acid metabolism — processes relevant for tumor growth and cancer progression. Interestingly, on the level of functional categories, we found significant enrichment for receptors in general (p-value 6.7 · 10^−14^, 1.9 fold enrichment) and specifically G-protein coupled receptors (p-value 1.7 · 10^−6^, 2.1 fold enrichment) as well as for substrate specific transmembrane transport (p-value 4.4 · 10^−9^, 2 fold enrichment), pointing to a possible involvement in signaling. On the component level, we found significant enrichments of the plasma membrane (p-value 2.2 · 10^−20^, 1.6 fold enrichment) and the extracellular region in general (p-value 4 · 10^−26^, 2.2 fold enrichment). The interesting enrichments on the process level include ion transport (p-value 1.3 · 10^−13^, 2 fold enrichment) and cell adhesion (p-value 1 · 10^−10^, 1.9 fold enrichment). All these results are plausible in the light of what is known about cancer biology and will be further investigated. The identified cancer-specific isoforms have a potential use as diagnostic marker or as possible drug targets.

### 3.2 Identification of SNVs Associated with Splicing Changes

After preprocessing and filtering, we have analyzed 11, 383 exon skip events as well as 3, 961 alternative 5’- and 5, 038 3’-end events. All events have been associated with 458, 266 variants. Since each event can represent the same transcript structure, these events are certainly not independent leading to exon skip events in 5, 623 genes, 3, 703 genes with alternative 3’-ends and 3, 278 genes showing alternative 5’ ends. After a very conservative correction (p-value *<*5 · 10^−9^ and 5% effect size), we find 251 polymorphic sQTLs of which 228 are *cis*-QTLs and 23 are *trans*-QTLs. Full break down in Table 1 and an overview of all sQTLs can be seen in Figure 2. It clearly demonstrates the generally higher power in detecting *cis*-QTLs due to the more direct nature of their effects and thus are easier to detect.

**Table 1.** Break down of sQTL associations. This table shows how many sQTLs with more than 10% effect size and p-value < 5 · 10^−7^ p-value and < 5 · 10^−9^ (Filtered) are found to be annotated in various functional databases. The top two rows sum to all 915 sQTLs detected and subsequent subsets are shown below.

| Category | Exon Skip (Filtered) | Alt 3' (Filtered) | Alt 5' (Filtered) |
| --- | --- | --- | --- |
| *cis* | 228(127) | 101(62) | 62(39) |
| *trans* | 312(12) | 104(8) | 108(3) |
| *cis* ClinVar | 2(1) | 3(2) | 0(0) |
| *trans* ClinVar | 2(0) | 0(0) | 1(0) |
| *cis* COSMIC | 18(11) | 8(4) | 1(0) |
| *trans* COSMIC | 6(1) | 1(0) | 2(0) |
| *cis* GWAS | 1(1) | 2(1) | 0(0) |
| *trans* GWAS | 1(0) | 0(0) | 0(0) |

**Fig. 2.**
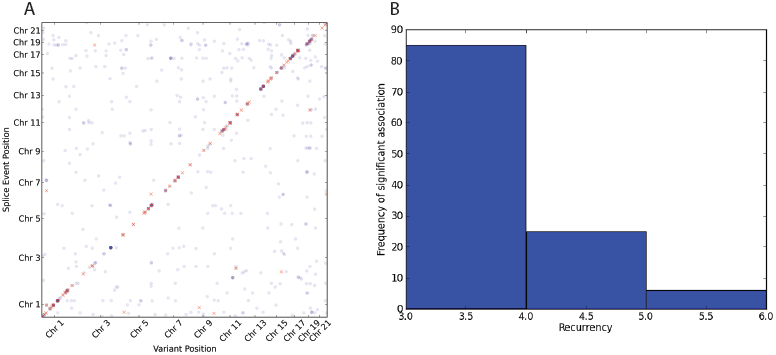
Summary plots of associations found in somatic and germline calls. *A*: Germline sQTL overview. Every dot represents a sQTL. Results have been filtered for 5% effect size. Red dots indicate sQTLs with p-value < 5 · 10^−9^ and blue dots indicate sQTLs with p-value < 5 · 10^−7^. *B*: Histogram indicating how many variants are found recurring (x-axis) in a given number of samples (y-axis).

#### 3.2.1 Associating Somatic Variants

We also considered a small set of 128 recurrent somatic mutations that are expected to be highly enriched with functional variants. Surprisingly, we found a large fraction of those somatic variants to be associated in *trans* and none in *cis* (p-value < 5 · 10^−4^).

Out of these associations, five are annotated COSMIC variants for which we have found none of them to be significant in our previous analysis on the tumor germline calls. While it could be reasoned that most somatic variants may have a functional effect, it is notable that a high fraction is significant after Bonferoni correction and that most of them are rare (see Figure 2 B). While we are confident in our statistical analysis, technical and biological validation will ultimately be able to separate false positive from biological meaningful results.

#### 3.2.2 ClinVar Annotated sQTL Suggest Functional Mechanisms

In an effort to establish links of interest between sQTLs and existing variants of interest we have compared our results to variants annotated in ClinVar. This analysis revealed that two polymorphic sites both associated (p-value < 2.6 · 10^−8^ and p-value < 1.1 · 10^−9^) with the same alternative 3’ splicing event within the Paraoxonase 2 gene have previously been associated with risk for coronary heart disease.^25^ We found another variant associated with three different alternative splice events in the ACP1 gene. While this variant is of benign clinical importance, we have confirmed previous mechanistic insights.^26^ Another polymorphic site in the SOD 2 gene is of clinical interest and is associated with an exon skip variation within the same gene. This variant is associated with increased risk of nephropathy in diabetics. This gene is of particular interests since variants in this gene have been associated with various diseases as in idiopathic cardiomyopathy (IDC), premature aging, sporadic motor neuron disease, and cancer (genecards source). A type 1 diabetes risk missense variant has also been associated with an alternate 3’-end in the OAS1 gene and it is known that variants influencing alternative splicing in this gene are of functional importance. It is notable that this region is also spanned by a long non-coding RNA (ENSG00000257452).^27^ We have also linked a risk variant for Myocardial infarction as well as an autoimmune disease susceptibility variant as sQTL variants with an exon skip event and an alternate 5’-event, respectively. This analysis demonstrates successfully how our sQTL analysis can confirm and suggest mechanistic insights into clinically and molecularly significant phenotypes.

#### 3.2.3 COSMIC Annotated sQTL

We identified 16 *cis*-sQTLs under very conservative thresholds (p-value < 5 · 10^−9^ and 0.25% effect size), which are also annotated in COSMIC suggesting their potential effect and involvement in cancer. Of particular interest are those variants annotated as sQTL in commonly mutated cancer genes.

Fig. 3 demonstrates an example where the most significant variant is annotated in COSMIC and shows a large difference in the splicing index across the different alleles in gene PMF1. This gene is known to be associated with bladder carcinoma and thus is of specific interest. While this somatic variant is rare, it overlaps a more common germline variant and thus did allow us to identify it as an sQTL in this study.

**Fig. 3.**
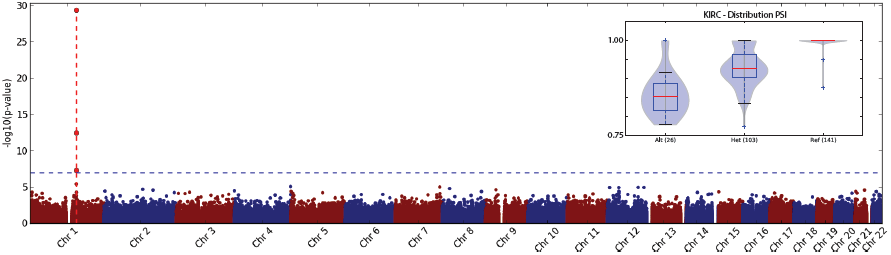
An example of a COSMIC annotated variant with alternative splicing. The dotted red line indicates the position of the alternative splice event. The figure on the top right shows a violin plot of PSI demonstrating the shift of distribution for each of the three possible genotype across all samples.

We found that the tumor necrosis factor related protein encoded by the C1QTNF3 gene harbors a recurrent COSMIC variant (COSM449566) which also appears to be a *cis* sQTL.This suggests potential functional effects with an effect size of ∼30.25%. Further analysis of such variants may be promising in understanding the effect of some commonly observed somatic variants.

#### 3.2.4 sQTL as Susceptibility loci for Cancer

A comparison of the set of sQTL with a known catalogue of genome-wide association study (GWAS) hits did yield four loci which have been linked to previous studies.^28^ Interestingly, three out of these loci are susceptibility loci for different types of cancers while the other one has been previously linked to multiple sclerosis. A variant introducing an alternate splice junction which then leads to a change of the alternate 3’-end of exon two and subsequently causing a truncation of that exon has previous been identified in two association studies. This variant is being highly associated with changes in serum magnesium levels (PMID) and it has also been identified as a susceptibility locus for gastric adenocarcinoma and esophageal squamous cell carcinoma. An sQTL in a mitochondrial carrier protein (SLC25) has recently been identified as a potential new susceptibility locus for testicular cancer^29^ which is of particular interest with respect to our finding of SLC enrichment. The well-studied BABAM1 gene appears to host an sQTL which is also an annotated COSMIC variant which is associated in several publications with hormone receptor-negative breast cancer and is also a susceptibility locus for ovarian cancer^30,31,32,33 34^ These results are encouraging and may be suggestive of alternative splicing patterns as underlying mechanism for cancer progression rather than as functional consequence.

## 4. Discussion

We have completed an extensive analysis on RNA-Seq samples originating from patients with Kidney Renal Clear Cell Carcinoma and present the first systematic sQTL analysis on kidney cancer to our knowledge. State-of-the-art methods are being used to systematically investigate splicing patterns in KIRC samples. We have developed a ranking mechanism to identify splicing events specific to tumor samples in comparison with normal samples. Our analysis demonstrates that we do not only find sQTL for tumor specific splicing events 1 B but are also able to identify functionally annotated variants providing potential new mechanistic insights.

Our analysis revealed a subset of genes that showed large differences in splicing events between tumor and normal samples. Although we tried to control for tissue-specific expression by taking the normal samples into account, this effect may still be partially confounded by cancer specific gene expression (see supplemental Fig. 1). While these genes may certainly be interesting with regards to being cancer markers, it is not absolutely clear whether altered functions associated with the observed events in these genes are driving tumor progression or are mere passenger events. However, considering that several top ranked genes have been previously linked to cancer metabolism and treatment outcome encourages further molecular analyses of these findings. TMEM176 may be a new interesting target due to the identification of tumor specific intron expression as well as the identification of sQTL associated with this event.

The analysis of the variants found in the matched whole exome sequencing data and their association with patterns of alternative splicing revealed various germline and COSMIC variants either directly causal or in linkage disequilibrium with causal variants. Some of the *cis*-sQTLs have previously been annotated as being susceptibility loci for cancer or other diseases and our analysis suggests a potential mechanistic involvement of splicing aberrations. We were further able to link various sQTLs to variants annotated in ClinVar suggesting splicing related mechanisms. In order to understand to what extent somatic mutations may be driving splicing changes, we have analyzed a small set of somatic variants identified by matching tumor normal pairs. After Bonferroni correction across the events and variants tested, most of the identified somatic variants remain significant which causes some concerns with regard to the amount of false positives in this set. Further analysis demonstrates that most of the associations are seen in less than four samples. While this might be an intrinsic property of somatic mutations, we suggest to explore also different approaches to take the lower frequency of somatic variants into account, specifically addressing false positive results. However, we are encouraged that many of the associations we found are indeed correct and meaningful in the context of cancer biology.

Our analysis is a step forward towards gaining further insight into the involvement of splicing patterns in cancer. It still remains to be seen to what extent alternative splicing patterns can create large changes in phenotypic outcome. Studies of natural population suggest that the effect of germline variants is generally small, however our results suggest that several germline and somatic variants may contribute towards functional changes. While we can confirm previously known alternative splicing events and the genetic markers driving these splicing changes, the functional role of many of these genes and changes in splicing patterns in cancer is unknown. We believe that new approaches and larger sample sizes are needed to gain further insight into the role of somatic variants. Our future work will involve addressing these issues and including samples of larger sizes from the TCGA project to gain more power to study the effect of rare somatic variants in cancer. This will involve the integration of rare-variant analysis approaches and integration of whole-genome data generated by the TCGA and ICGC project.

## Acknowledgements

We gratefully acknowledge discussions with Ari Hakimi, Ed Reznik and Chris Sander. This work is funded by the Sloan Kettering Institute (to G.R.) and was relying on the Beckman genomic data storage facility (Geoffrey Beene grant to G.R. & N.S.). Partial funding under NIH grant 1R01CA176785-01A1.

## Supplemental materials

Additional material including a detailed method description can be found under the following url: http://www.raetschlab.org/suppl/kirc-splicing

